# Weight of evidence of Y-STR matches computed with the discrete Laplace method: Impact of adding a suspect’s profile to a reference database

**DOI:** 10.1101/2022.08.25.505269

**Authors:** Mikkel Meyer Andersen, Poul Svante Eriksen, Niels Morling

## Abstract

The discrete Laplace method is recommended by multiple parties (including the International Society of Forensic Genetics, ISFG) to estimate the weight of evidence in criminal cases when a suspect’s Y-STR profile matches the crime scene Y-STR profile. Unfortunately, modelling the distribution Y-STR profiles in the database is time-consuming and requires expert knowledge. When the suspect’s Y-STR profile is added to the database, as would be the protocol in many cases, the discrete Laplace model must be recomputed. We found that the likelihood ratios with and without adding the suspect’s Y-STR profile were almost identical with 1,000 or more Y-STR profiles in the database for Y-STR profiles with 8, 12, and 17 loci. Thus, likelihood ratio calculations can be performed in seconds if a an established discrete Laplace model based on at least 1,000 Y-STR profiles is used. A match in a database with 17 Y-STR loci from at least 1,000 male individuals results in a likelihood ratio above 10,000 in approximately 94% of the cases, and above 100,000 in approximately 82% of the cases. We offer a freely available IT tool for estimating the discrete Laplace model of the STR profiles in a database and the likelihood ratio.

**Highlights:**

- The discrete Laplace method is suitable for estimating the weight of evidence of matches with 17 Y-STRs.
- *LR*s based on the discrete Laplace method are 10-100 times higher (in median) than those based on Brenner’s *κ* method.
- A database with 17 STRs from at least 1,000 males gives *LR*s of above 10,000 in approximately 94% of the cases and above 100,000 in approximately 82% of the cases with the discrete Laplace method.
- The weight of evidence of a matching Y-STR profile is computed within seconds and easily documented when a precomputed discrete Laplace model is available (an IT tool is provided).
- 50% of all Yfiler Plus matches are between male relatives within a genetic distance of five meioses.

## 1. Introduction

Using Y chromosomal short tandem repeat (STR) profiles (Y-STR profiles) in forensic genetic casework includes estimation of the evidential weight of a match between a suspect’s Y-STR profile and a crime scene Y-STR profile. Several methods for estimating the weight of evidence of Y-STR profile matches have been suggested [1–9]. A review of the topic is found in [1]. Here, we focus on the discrete Laplace method [2] recommended by the DNA commission of the International Society of Forensic Genetics (ISFG) [10], the Y-chromosomal short tandem repeat haplotype reference database (YHRD) [11], and Andersen and Balding (2021) [1] (provided that the Y-profiles are suitably slowly mutating to ensure the matching individuals are found in the entire population and not only among the suspect’s close paternal relatives).

The discrete Laplace method requires a reasonably sized Y-STR profile reference database with not too many Y-STR loci due to inherent statistical issues (e.g. the curse of dimensionality). There are various practises of including two, one, or zero copies of the Y-STR profile of the suspect in the reference database for estimating the Y-STR profile probability. The argument for adding two Y-STR profile copies is that the suspect and donor are two different individuals under the alternative hypothesis in the likelihood ratio calculation (the denominator). Hence, both the suspect’s and donor’s profiles are added to the reference database. This may be acceptable for large autosomal STR reference databases since adding two autosomal STR allele copies does not change the estimated allele frequencies much. However, many Y-STR profile databases are small, and adding two copies of the suspect’s Y-STR profile may inflate the estimate of the Y-STR profile probability. Furthermore, the Y-STR profiles are not independent samples because the suspect’s profile is only considered because it matches the donor’s profile. Therefore, we do not agree to add two copies of the suspect’s Y-STR profile to the reference database for estimating the Y-STR profile probability. However, we think that adding the suspect’s Y-STR profile to the reference database is sensible, as also done by, e.g. [2, 3].

We studied the effects of adding and not adding the suspect’s Y-STR profile to the reference data. The reason for not adding a copy of the suspect’s Y-STR profile is that if a new Y-STR profile is added to the reference data, the discrete Laplace model must be recomputed. The recomputation takes several minutes to several hours with conventional computers followed by an expert’s model inspection and sanity checks. For routine criminal casework, calculating the evidential weight should be fast, easy, and robust. Thus, discrete Laplace model computations with the addition of the Y-STR profile of the matching suspect have prevented the practical use of the method because many Y-STR databases are relatively small. However, if sufficiently large Y-STR profile databases, e.g., national databases, are established, a single discrete Laplace model can be used for all database computations. The remaining question is how large the Y-STR profile database should be so that there, for practical purposes, is no difference between the weight of evidence calculated with and without adding the matching suspect’s Y-STR profile when using the discrete Laplace method.

Using Y-STR profile probability estimates as match probabilities is only acceptable if the Y-STR profiles are slowly mutating to ensure that the matching individuals are found in the entire population and not only in the suspect’s close paternal relatives, as discussed by [1, 7]. Hence, we focus on Y-STR profiles with overall mutation rates, i.e., mutation at one or more loci in a Y profile, lower than 5% [1]. Finally, we briefly discuss the issue with close male relatives of the matching suspect using the data from the simulation study by Andersen and Balding (2017) [7].

## 2. Materials and Method

### 2.1. Data and data analysis

Data were analysed using R [12] version 4.1.3 with the packages `disclapmix` version 1.7.4 [13] and `tidyverse` [14]. All code for running the analyses in this paper is available at [15].

In [16], 19,630 PPY23 Y-STR profiles from the entire world are available. We used data of individuals of European ethnicity living in Northern and Central Europe, excluding Finns, who are genetically different from other Northern Europeans. We excluded DYS385a/b due to duplications and substracted DYS389I from DYS389II as DYS389II measures the entire length of DYS389, including the DYS389I part. There were 25 Y-STR profiles with duplications at DYS19, DYS389II, DYS439, DYS448, DYS481, DYS533, DYS570, DYS576, and DYS635. These samples (not loci) were excluded. This resulted in a full reference database, *D*^0^, of size 5,823 individuals with data from 21 Y-STR loci (Table 1).

**Table 1:**
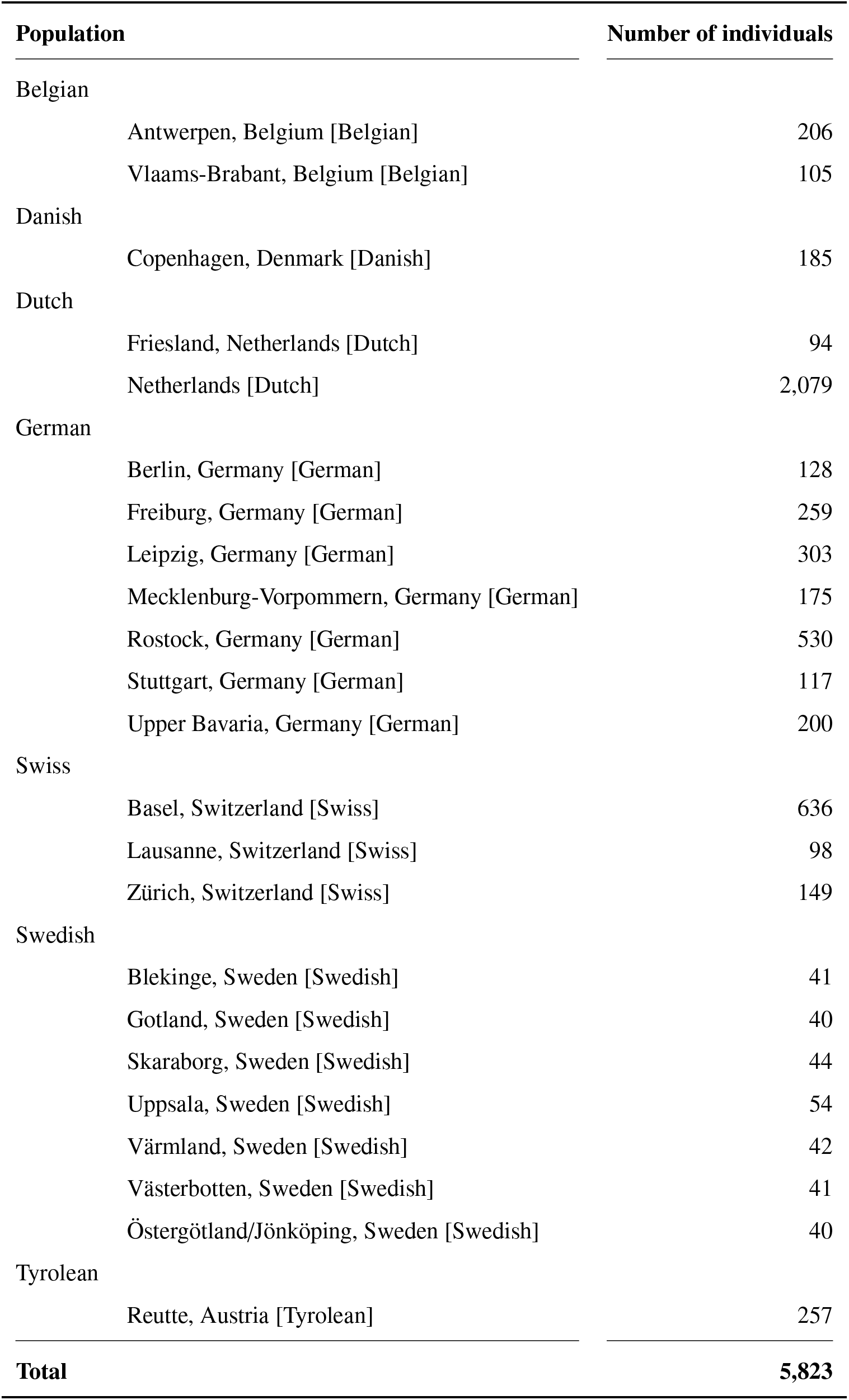
Y-STR population data used for the Y-STR reference database, *D*^0^.

| Population | Number of individuals |
| --- | --- |
| Belgian |  |
| Antwerpen, Belgium [Belgian] | 206 |
| Vlaams-Brabant, Belgium [Belgian] | 105 |
| Danish |  |
| Copenhagen, Denmark [Danish] | 185 |
| Dutch |  |
| Friesland, Netherlands [Dutch] | 94 |
| Netherlands [Dutch] | 2,079 |
| German |  |
| Berlin, Germany [German] | 128 |
| Freiburg, Germany [German] | 259 |
| Leipzig, Germany [German] | 303 |
| Mecklenburg-Vorpommern, Germany [German] | 175 |
| Rostock, Germany [German] | 530 |
| Stuttgart, Germany [German] | 117 |
| Upper Bavaria, Germany [German] | 200 |
| Swiss |  |
| Basel, Switzerland [Swiss] | 636 |
| Lausanne, Switzerland [Swiss] | 98 |
| Zürich, Switzerland [Swiss] | 149 |
| Swedish |  |
| Blekinge, Sweden [Swedish] | 41 |
| Gotland, Sweden [Swedish] | 40 |
| Skaraborg, Sweden [Swedish] | 44 |
| Uppsala, Sweden [Swedish] | 54 |
| Värmland, Sweden [Swedish] | 42 |
| Västerbotten, Sweden [Swedish] | 41 |
| Östergötland/Jönköping, Sweden [Swedish] | 40 |
| Tyrolean |  |
| Reutte, Austria [Tyrolean] | 257 |
| **Total** | **5,823** |

We made three combinations of Y-STR loci, each with a combined Y-STR profile mutation rate just below 5%, based on the mutation rates published at www.yhrd.org [11], accessed on October 5, 2021. The Y-STR loci combinations are listed in Table 2. We refer to them as “kits” for brevity.

**Table 2:** Combinations of Y-STRs with combined mutation rates below 5% as suggested by [1]. Data based on the mutation rates published at www.yhrd.org [11], accessed on July 15, 2022.

|  | 8 loci | 12 loci | 17 loci |
| --- | --- | --- | --- |
| **Overall mutation rate** | 0.0498 | 0.049 | 0.0462 |
| **Mutation rate range ($\times 10^{-3}$)** | 4.24 - 12.12 | 1.32 - 12.12 | 0.35 - 6.60 |
| **Loci**<br>**(DYS prefix removed)** | 576, 570, 458, 439,<br>389II, 481, 456, 635 | 576, 570, 439, 481,<br>456, 533, YGATA4,<br>391, 19, 390, 448,<br>437 | 438, 392, 393, 437,<br>448, 390, 19, 391,<br>389I, YGATAH4,<br>533, 635, 456, 481,<br>389II, 439, 458 |
| **All loci in PowerPlex Y23** | Yes | Yes | Yes |
| **All loci in Yfiler Plus** | Yes | Yes | Yes |
| **Comment** | Loci with the highest mutation rates | Only markers in Yfiler Plus, removing DYS389I/II, DYS458, and DYS635 (fairly high mutation rates) to allow for more slowly mutating loci | Loci in Yfiler Plus with lowest mutation rate |

Following [17], we calculated how much information each of the three kits contained compared to all 21 Y-STR loci. This was done by the uncertainty coefficient, *C* = *I*(*X*; *Y*)*/H*(*X*), where *Y* is the Y-STR loci in a kit, *X* is the remaining Y-STR loci out of the 21 ones not in *X* (such that *X* and *Y* together are all 21 Y-STR loci), *H*(*X*) the Shannon entropy [18], and *I*(*X*; *Y*) the mutual information [18]. Note, that the uncertainty coefficient, *C*, can take values from 0 (if *X* and *Y* are independent and does not provide information about each other) to 1 (when knowing *Y* leaves no uncertainty about the value of *X*). This was referred to as “Percentage of overall entropy explained by seven markers” (actually 8, 12, or 17 markers) by [17].

### 2.2. Reference databases

From the full reference database, *D*^0^, of size 5,823, we drew a reference database, *D*^+^, of size *n*. This was done 200 times for each of the following 16 values of *n*: 101, 201, 301, 401, 501, 601, 701, 801, 901, 1,001, 1,501, 2,001, 2,501, 3,001, 4,001 and 5,001. We drew the profiles from the full reference database (size 5,823) without replacement to reflect a sequential sample selection as the full reference database of size 5,823 does not consist of all possible Y-STR profiles worldwide. In each of the 16 × 200 = 3,200 reference databases, we then chose 5 “cases” for each database by randomly withdrawing a Y-STR profile from *D*^+^, resulting in a reference database, *D*^−^, of size *n* − 1. Thus, there were 16 × 200 × 5 = 16,000 “cases”, each analysed with the three kits in Table 2. The “cases” were used to analyse if a discrete Laplace model based on the database *D*^−^ gave similar Y-STR profile probabilities as one based on *D*^+^.

Let 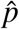 denote the estimated Y-STR profile probability such that 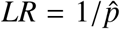 (with the cautions mentioned by [1] in regards to, e.g., close paternal relationship between the suspect and the true donor). Denote by

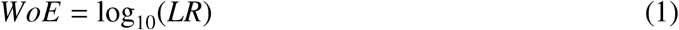

the weight of evidence in bans. Introduce *WoE*^−^ and *WoE*^+^ to refer to whether the computations were performed on the reference database without adding the suspect’s profile (size *n* −1) or with the suspect’s Y-STR profile added (of size *n*), respectively.

We included Brenner’s *κ* method [3] where relevant. Note that some of the databases of size 101, 201, and 301 contained Y-STR profiles that were all different, and the Brenner’s *κ* gives an *WoE*^+^ of infinity. Those “cases” were excluded for comparison with results obtained with Brenner’s *κ* method.

### 2.3. Discrete Laplace modelling

For each “case”, we estimated discrete Laplace models for 1 to *k* clusters ensuring that *k* was at least five clusters larger than the best model (lowest marginal Bayesian Information Criterion, BIC, value [2, 13]) with the `disclapmix_adaptive` function in the `disclapmix` package [13] (refer to the vignettes for examples; they are available in the package or directly at https://mikldk.github.io/disclapmix/articles/). We chose the following ways of using the discrete Laplace models to estimate a Y-STR profile probability, 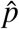:

- “Best”: The model with the lowest marginal BIC value was used to predict the Y-STR profile probability.
- “Top-5 weighted”: The five models with the lowest marginal BIC values were used to calculate the weighted Y-STR profile probability; the weights were the normalised reciprocal BIC values so the sum of the weights was 1.
- “Top-5 minimal”: The maximal estimated Y-STR profile probability of the Y-STR profile in question among the top-5 models, which results in the minimal *LR* as 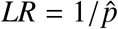.

### 2.4. Effect of adding the suspect’s matching Y-STR profile to the database

Let 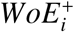 be the log_10_ of the *LR* of the *i*^th^ Y-STR profile based on *D*^+^. Similarly, let 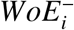 be the log_10_ of the *LR* of the *i*^th^ Y-STR profile based on *D*^−^. Denote by

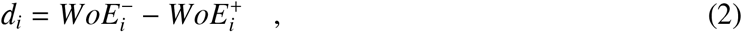

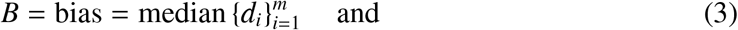

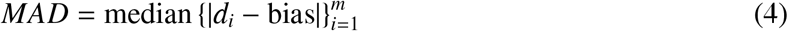

the deviation (*d*), the bias (*B*), and the median of the absolute deviations from the bias (*MAD*) between the logarithms of the *LR*s without and with adding the suspect’s matching Y-STR profile. Thus, *d* measures the *WoE* differences between adding and not adding the Y-STR profile. If *d*, e.g., is 2, the *LR* is 100 times larger without than with adding the suspect’s matching Y-STR profile. If *d*, e.g., is −2, the *LR* is 100 times larger with than without adding the suspect’s matching Y-STR profile, but *d* will usually not be negative, as we will see below. The bias is the median of the differences, i.e., an estimate of the expected difference between *WoE* with and without adding the Y-STR profile in question. We calculated the bias and *MAD* for each kit and database size, i.e., each *MAD* value was based on *m* = 200 × 5 = 1,000 “cases”.

## 3. Results

### 3.1. Proportion of information and numbers of Y-STR loci

Polymorphism increases when more and more Y-STR loci are examined due to the haplotypic nature of the Y chromosome. The information of, e.g., an Y-STR profile with 12 Y-STR loci contains more than 95% of the information contained in a Y-STR profile with 21 loci (Table 3) estimated by the uncertainty coefficient, *C* [17].

**Table 3:**
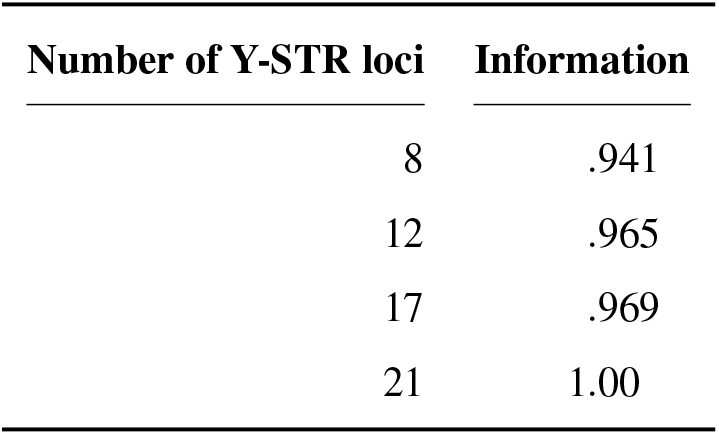
Percentage of information contained in three Y-STR loci subsets (Table 2) of 21 Y-STR loci. The information percentage was calculated as the uncertainty coefficient, *C*, that can take values from 0 to 1 (termed ‘percentage of overall entropy’ by [17]).

| Number of Y-STR loci | Information |
| --- | --- |
| 8 | .941 |
| 12 | .965 |
| 17 | .969 |
| 21 | 1.00 |

### 3.2. Proportion of Y-STR profile singletons

Very few males share Y-STR profiles with more than a handful of STR loci, and Y-STR profiles are mainly shared among close male relatives [7]. Many Y-STR profiles are not represented in even large Y-STR databases, and most Y-STR profiles are represented only once, called singletons. Fig. 1 shows the singleton proportions with 8, 12, and 17 Y-STR loci and selected database sizes from 100 to 5,000 Y-STR profiles.

**Fig. 1:**
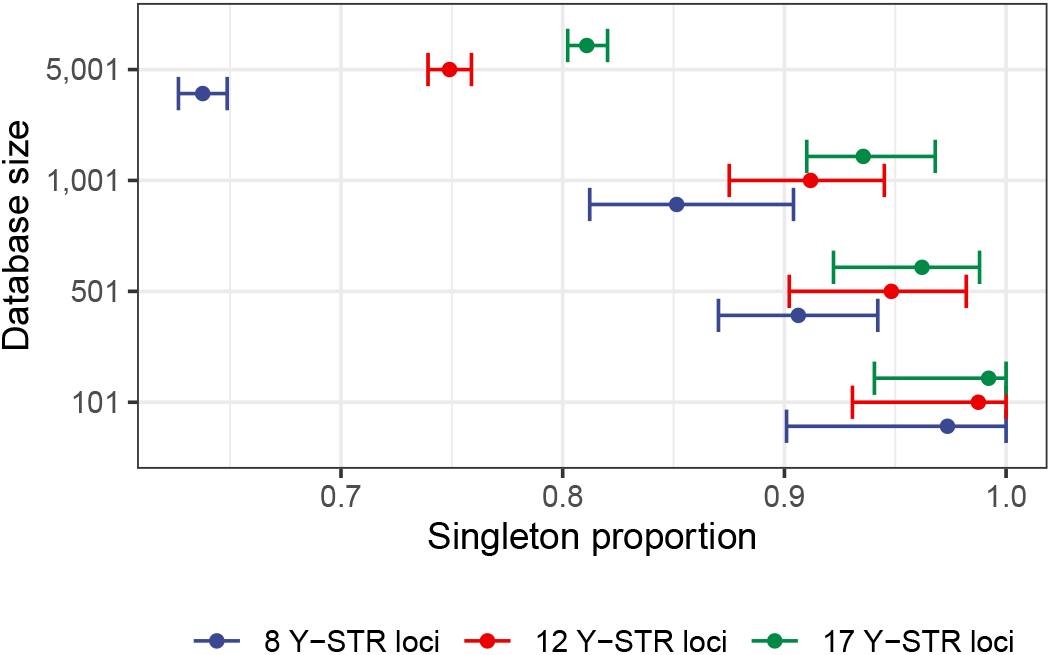
Singleton proportions for Y-STR database sizes 101, 501, 1,001, and 5,001. The points are the mean singleton proportions, and the intervals show the observed minimum and maximum singleton proportions.

### 3.3. Weight of evidence as a function of the size of a reference database

Fig. 2 compares the median *WoE*^+^ for different sizes of the reference database. The *WoE*^+^s based on the discrete Laplace method seem independent of database size but vary slightly due to random sampling of the reference databases, whereas the *WoE*^+^ based on Brenner’s *κ* systematically increases with the size of the reference database. Generally, the median *WoE*^+^s based on the discrete Laplace method are 1-2 bans higher than those based on Brenner’s *κ* method. Transformed to the *LR* scale, this corresponds to the median *LR*s based on the discrete Laplace method are 10-100 times higher than those based on Brenner’s *κ* method.

**Fig. 2:**
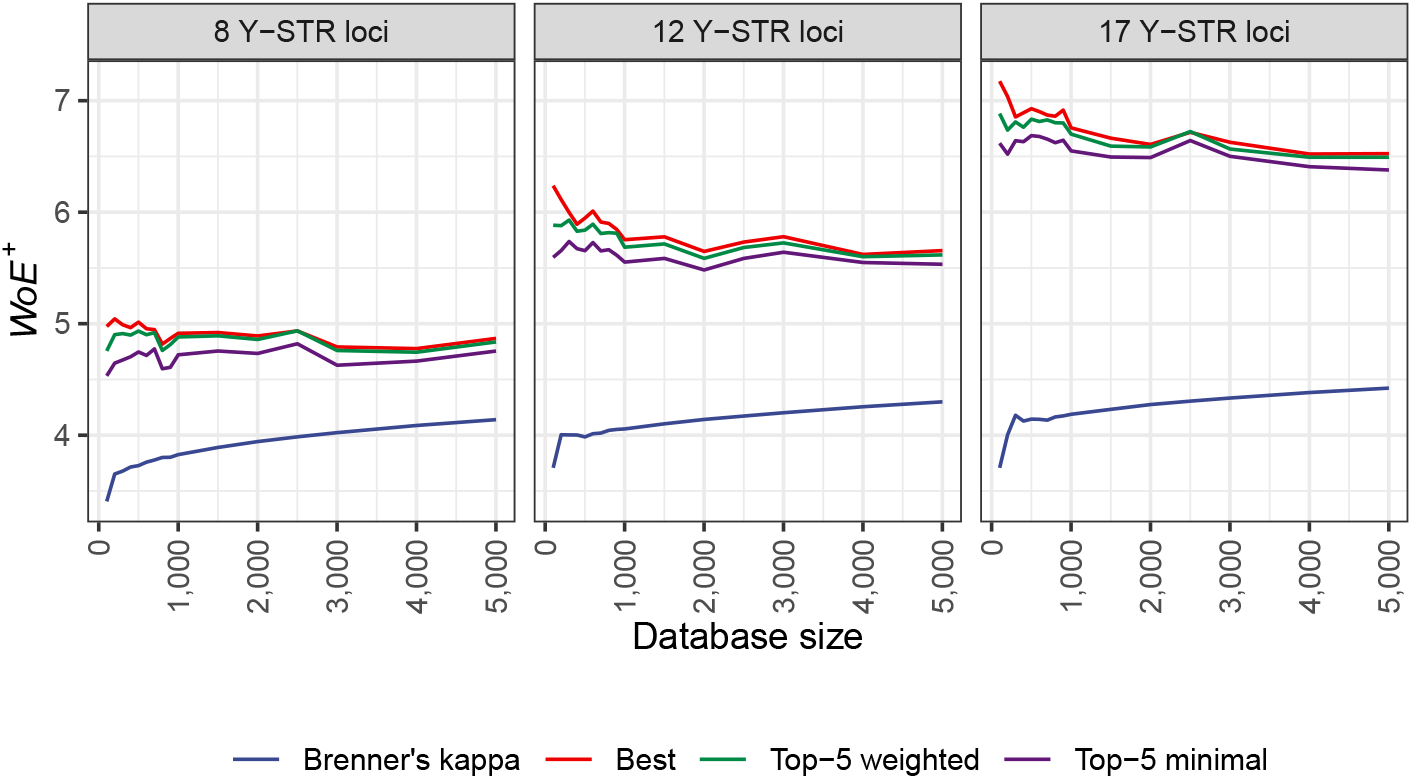
Comparison of the median *WoE*^+^ based on the extended database, *D*^+^, between the different methods. The methods Best, Top-5 weighted, and Top-5 minimal all refer to the discrete Laplace method with minor differences in the use of the models, as described in Sec. 2.3.

### 3.4. Weight of evidence with and without adding the suspect’s Y-STR profile

Fig. 3 compares the *WoE*s with and without adding the suspect’s Y-STR profile obtained with the best discrete Laplace model (lowest marginal BIC value). The correlation between the *WoE*s was high, although the *WoE*^+^ tended to be slightly lower than the *WoE*^−^. The more loci, the more variability around the identity line (at which the estimated probability is the same whether or not the Y-STR profile is added).

**Fig. 3:**
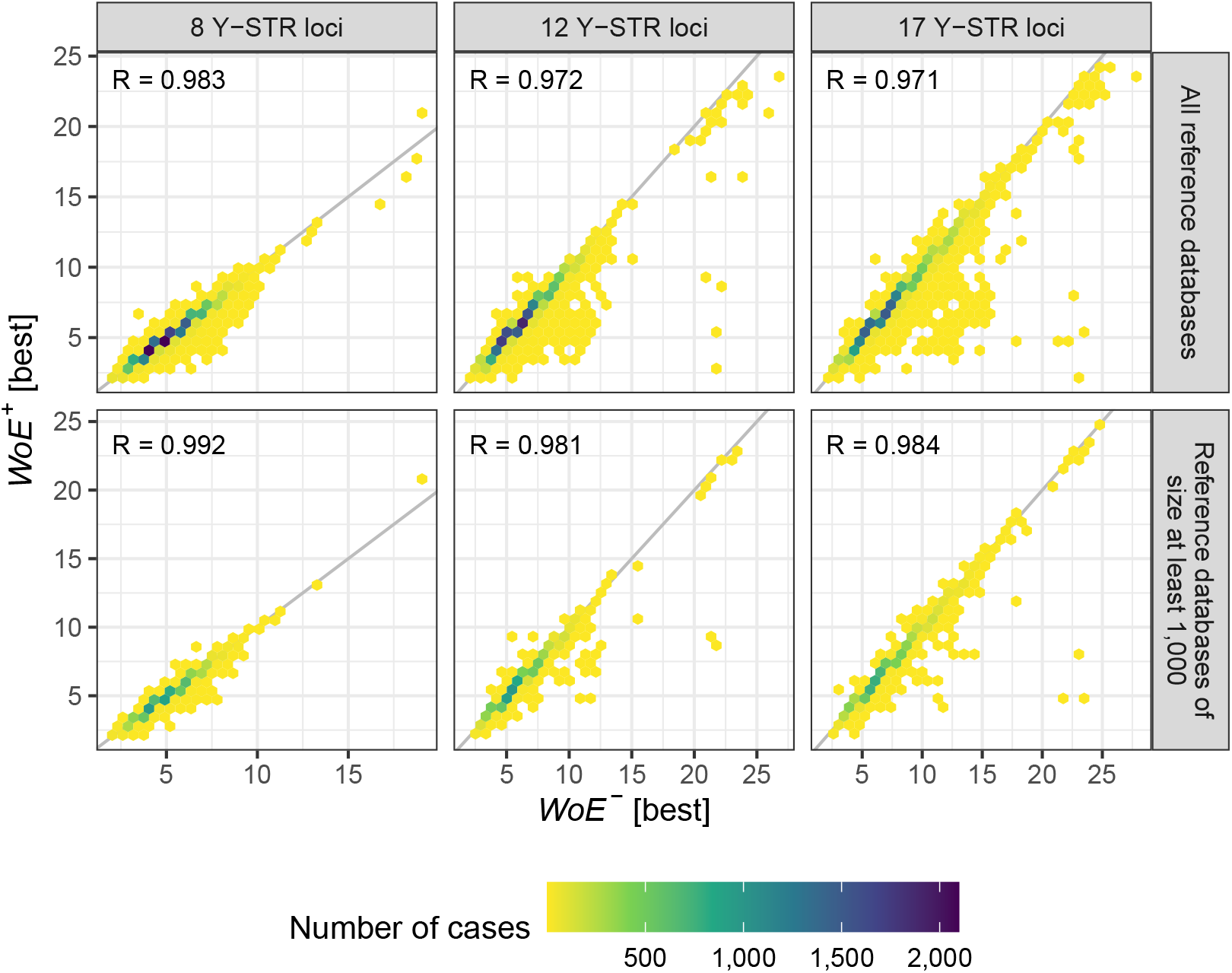
Comparison of the *WoE*s of the best discrete Laplace model (lowest marginal BIC value) obtained with and without adding the suspect’s Y-STR profile (*WoE*^−^ [best] versus *WoE*^+^ [best]) for all 16,000 “cases” for each kit, i.e., regardless of the database size in the first row, and only those of sizes above 1,000 in the second row. The identity line shows where the estimated probabilities are equal with and without adding the suspect’s matching Y-STR profile. *R* denotes Pearson’s correlation coefficient.

### 3.5. Weight of evidence of Y-STR profiles

Fig. 4 shows the percentages of cases above various *WoE* thresholds, and Table 4 shows the values of selected thresholds.

**Fig. 4:**
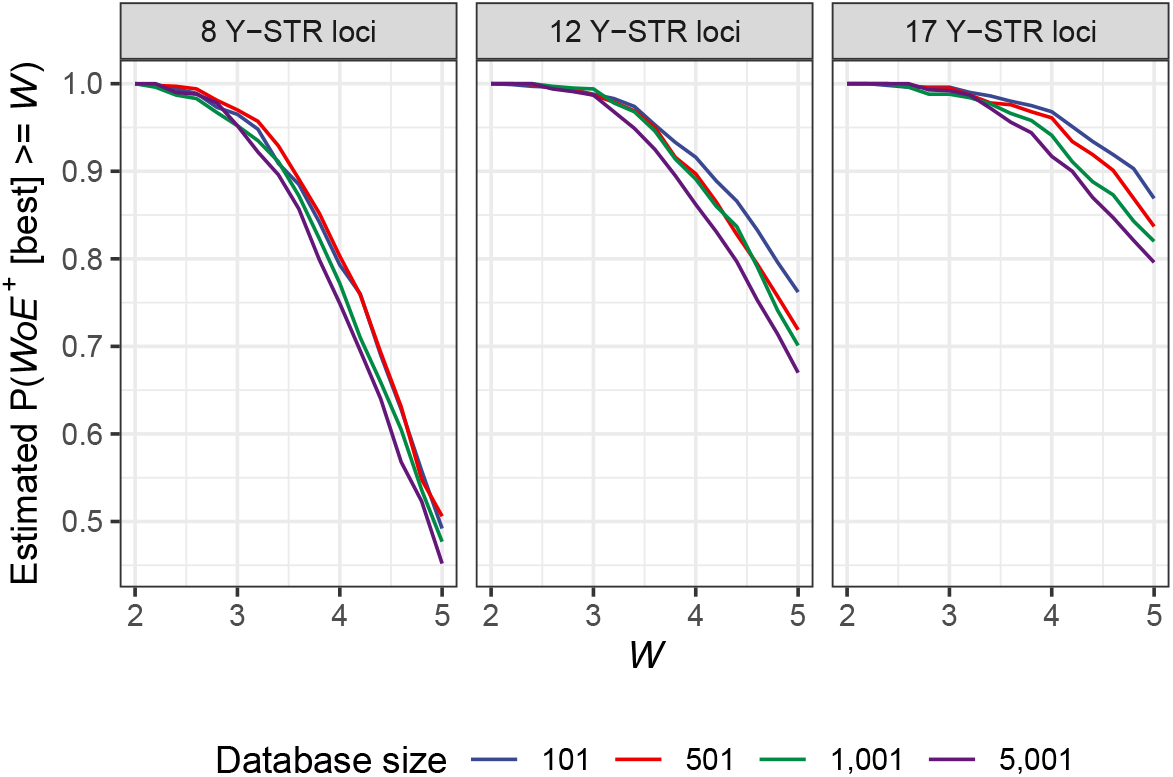
Percentages of cases above a certain *WoE*^+^. *WoE*^+^ is the log_10_ of the *LR* based on the extended reference database, *D*^+^, with the suspect’s Y-STR profile added. Data based on the best discrete Laplace model (lowest marginal BIC value). See Table 4 for the selected thresholds.

**Table 4:** Weight of evidence (*LR*) based on the discrete Laplace method with 8, 12, and 17 Y-STRs. Percentages of “cases” above *WoE*^+^ of above 3, 4, and 5 corresponding to *LR*s above 1,000, 10,000, and 100,000, respectively. *WoE*^+^ is the log_10_ of the *LR* based on the extended reference database, *D*^+^, with the suspect’s Y-STR profile added. Data based on the best discrete Laplace model (lowest marginal BIC value). See Fig. 4 for a visualisation with more threshold values.

| $WoE^+$ threshold | Threshold on $LR$ scale | 8 loci | 12 loci | 17 loci |
| --- | --- | --- | --- | --- |
| 3 | 1,000 | 95% | 99% | 98% |
| 4 | 10,000 | 77% | 89% | 94% |
| 5 | 100,000 | 47% | 70% | 82% |

### 3.6. Bias and variability of the weight of evidence

Fig. 5 shows the difference in the *WoE* measured by the *B* (bias) defined in Eq. (3) and *MAD* (median of the absolute deviations of *WoE* from the bias) defined in Eq. (4) with and without adding the suspect’s Y-STR profile to the database.

**Fig. 5:**
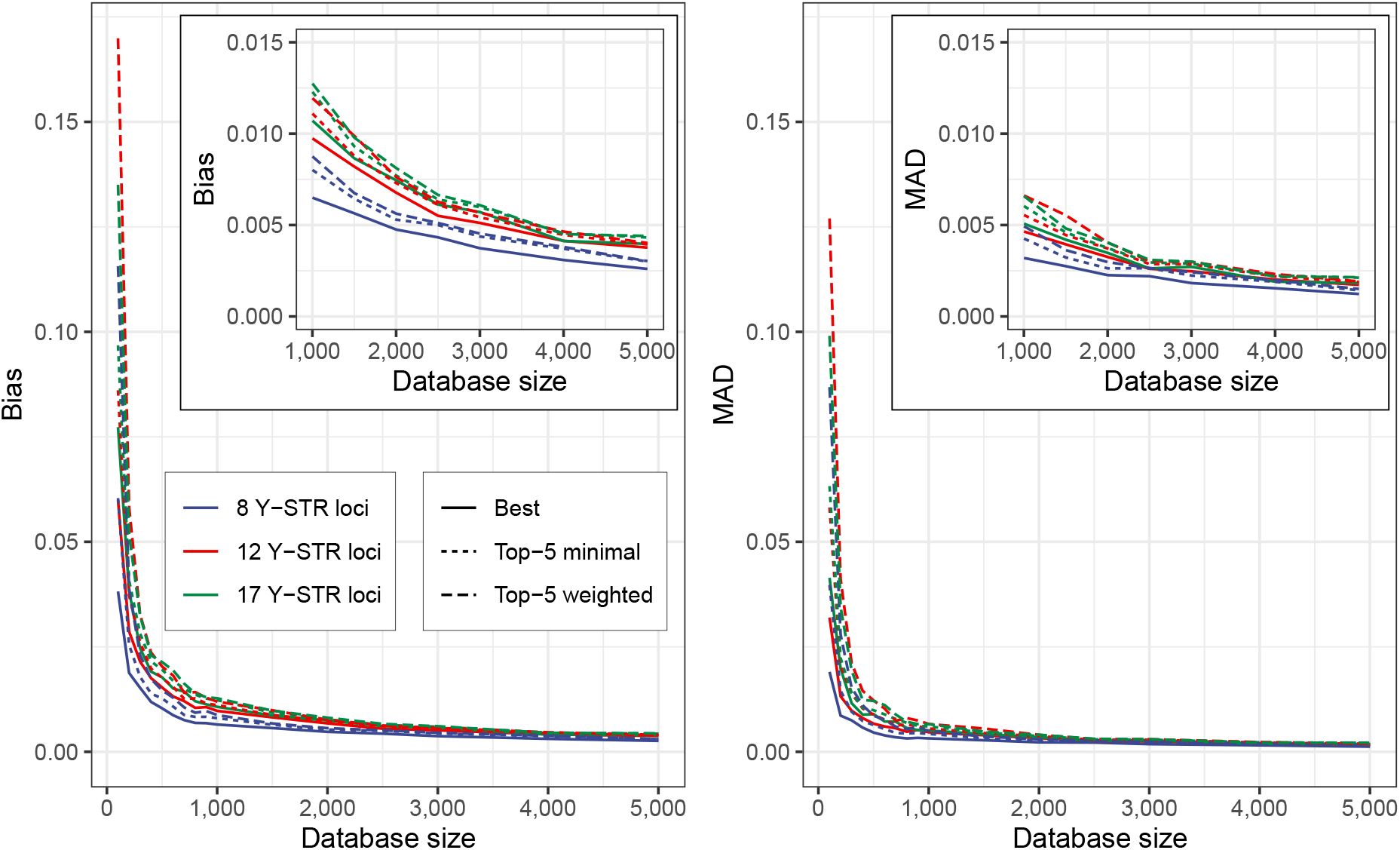
Differences in the *WoE* measured by the *B* (bias) defined in Eq. (3) (left hand side figure) and *MAD* (median of the absolute deviations of *WoE* from the bias) defined in Eq. (4) (right hand side figure) with and without adding the suspect’s Y-STR profile to the database. The inserted figures show the results with database sizes from 1,000 to 5,000 Y-STR profiles.

Fig. 6 shows the quantiles of the empirical distribution of differences in the *WoE* measured by *d* (deviations) defined in Eq. (2) for the best discrete Laplace model (lowest marginal BIC value) with and without adding the suspect’s matching Y-STR profile to the database. The larger the database size, the less the *WoE* changes when adding the suspect’s Y-STR profile.

**Fig. 6:**
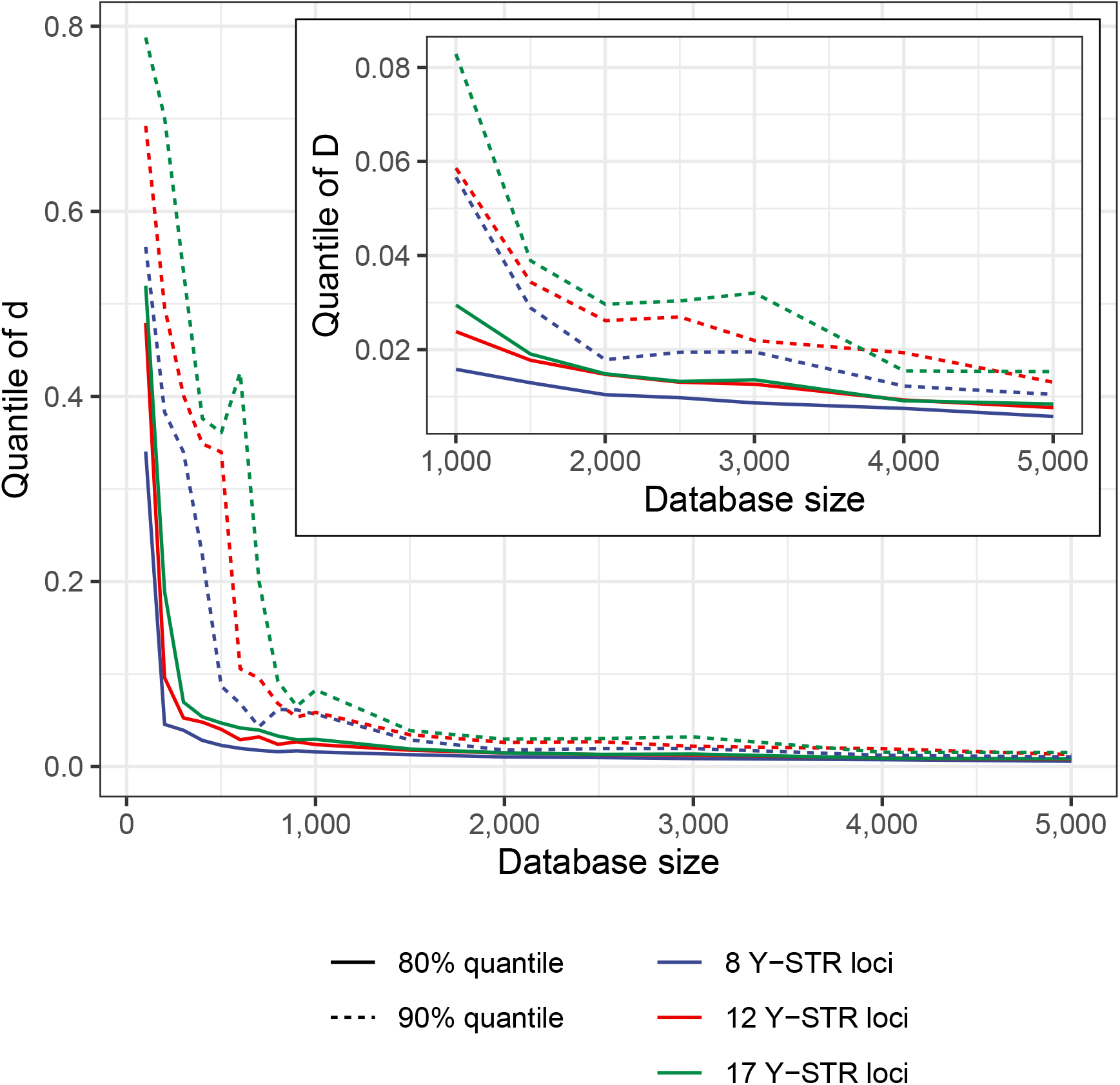
Empirical quantiles of differences in the *WoE* measured by *d* (deviations of *WoE*) defined in Eq. (2) for the best discrete Laplace model (lowest marginal BIC value) with and without adding the suspect’s Y-STR profile to the database.

### 3.7. Proportion of Y-STR matches among close relatives

The `malan` method [7] describes the distribution of the number of males sharing the same Y-STR profile. Unfortunately, the papers on the `malan` method do not include any analysis of the proportion of matches within various meiotic distances. Fig. 7 shows the data analysis of the Yfiler Plus kit (25 Y-STR loci counting duplicated loci only once). Approximately 50% of the matching males are related within a genetic distance of five Y-STR chromosomal meioses.

**Fig. 7:**
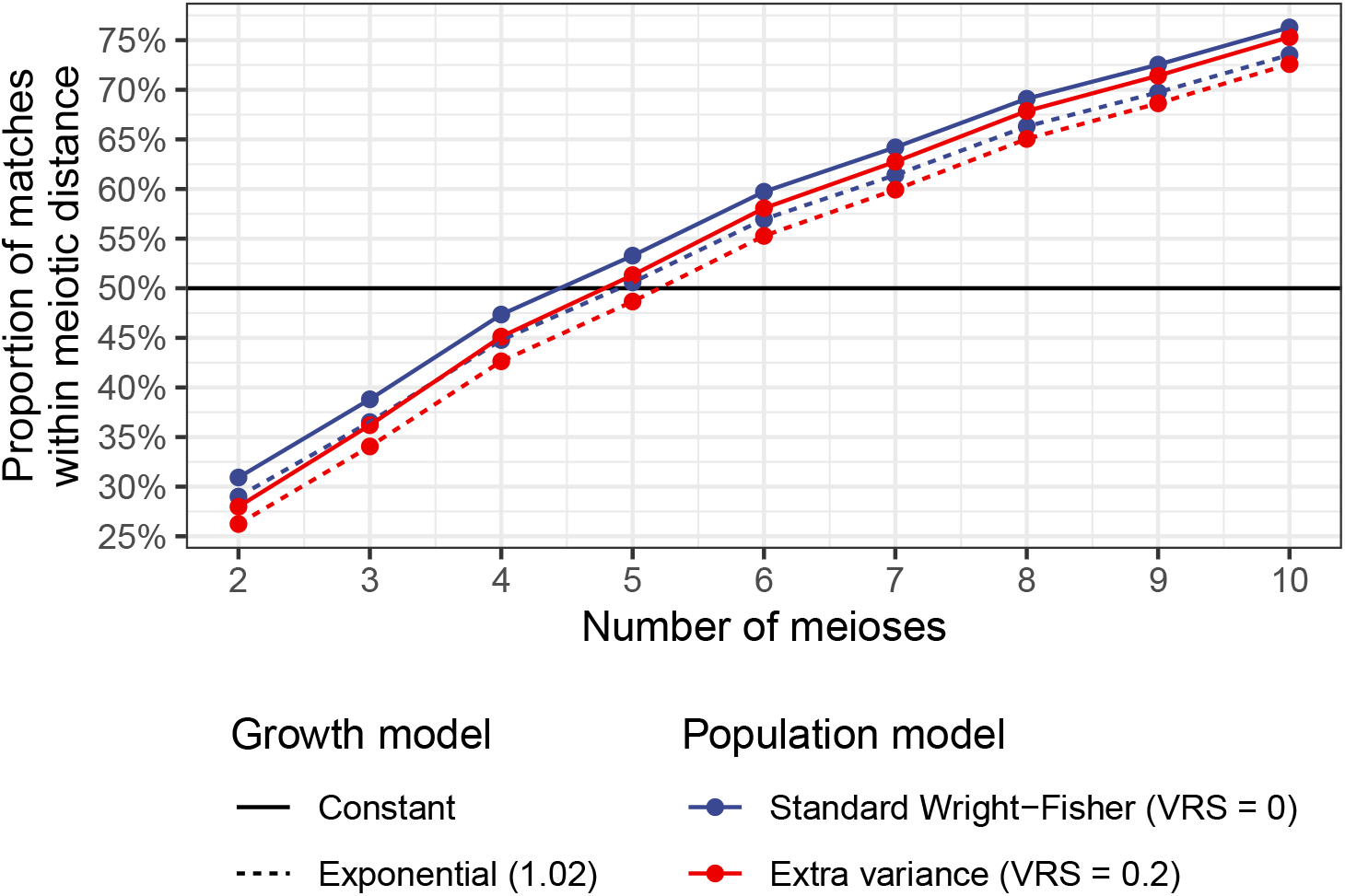
Proportion of matches for Yfiler Plus (25 Y-STR loci counting loci with duplications only once) within various meiotic distances based on simulated data from [7].

## 4. Discussion

### 4.1. Estimation of the weight of evidence of matching Y-STR profiles

Typing Y chromosome markers, including Y-STRs, are valuable tools in forensic genetics. Y-STR typing is mainly used in criminal cases and particular relationship cases. The polymorphism of Y-STRs is high and allows discrimination between unrelated males [7]. However, the very high proportion of close male relatives with shared Y-STR profiles makes it difficult to discriminate between close male relatives, even with more than 20 Y-STRs [7]. Y-STRs with high mutation rates have been selected to increase the discriminatory power among related and unrelated males [19].

Estimating the weight of evidence should, when possible, be based on the likelihood principle, both in relationship testing [20] and criminal cases [21]. In criminal cases, a relevant likelihood ratio should be calculated. In cases without close male relatives to the suspect, the weight of evidence can be expressed as

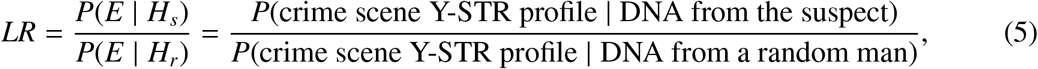

where *LR* is the likelihood ratio, *P* the probability, *E* the crime scene Y-STR profile, *H*_*s*_ the hypothesis that the DNA is from a suspect with a Y-STR profile matching the DNA evidence from, e.g., a crime scene, and *H*_*r*_ the hypothesis that the DNA from the crime scene is from a random man. Besides the problem related to a possible unknown close relationship [1], the extreme rarity of the various Y-STR profiles in most populations makes estimating the probability of Y-STR profiles difficult. The population Y-STR profile probabilities are based on observations in a database with a certain number of males. Most Y-STR profiles in a database are only represented once, i.e., singletons, and many Y-STR profiles are not represented in a database. Thus, the population probabilities of the Y-STR profiles are overestimated if a naïve calculation method like (*x* +1)*/*(*n* +1) or (*x* +2)*/*(*n* +2) is used (*x* is the number of times a profile appears in a reference database of size *n*). This leads to a considerable underestimation of the *LR*. Brenner’s *κ* method [3] compensates, to some degree, for the over-estimation of the population probability of rare Y-STR profiles.

### 4.2. The discrete Laplace method

We developed the discrete Laplace method to estimate the population probabilities of Y-STRs [2, 13]. The method is recommended by the DNA commission of the International Society of Forensic Genetics (ISFG) [10], the Y-chromosomal short tandem repeat haplotype reference database (YHRD) [11], and Andersen and Balding (2021) [1] (provided that the Y-profiles are suitably slowly mutating). In a criminal case, it is sensible to add the suspect’s matching Y-STR profile to the database. This means that the discrete Laplace model must be recomputed for all new cases if the database includes less than 1,000 Y-STRs making the practical use of the method slow. The computations involved in creating the discrete Laplace model based on the Y-STR profiles in a database are time-consuming (hours) due to computations, model inspection, and sanity checks by an expert. When the discrete Laplace model for a data set is established, the *LR* can be calculated in seconds, and the computations are easily documented. The importance of adding the suspect’s Y-STR profile is reduced by an increasing number of Y-STR profiles in the database. We investigated the practical effects of adding the suspect’s Y-STR profile to reference databases of various sizes before estimating the discrete Laplace model. We restricted the combined Y-STR profile mutation rate to a maximum of 5% [1] to ensure the matching individuals are found in the entire population – not only among the suspect’s close paternal relatives.

Since there is no published, sufficiently large Y-STR database from a single homogeneous population, we constructed a reference database, *D*^0^, of size 5,823 based on multiple smaller regional samples [16]. The ethnic and geographic composition does not reflect the proportions of the populations in North/Central Europe. Thus, it was not intended to make a North/Central European reference database. If a single homogeneous database were used, we assume that the *B* (bias) and *MAD* values would be decreased (Fig. 5).

The bias defined in Eq. (3) was positive (Fig. 5) such that the *WoE* and *LR* were larger when using the reference database without the profile in question (*D*^−^) than when using the reference database with the profile in question (*D*^+^). For reference databases of sizes of at least 1,000, the bias was smaller than 0.015 on a *WoE* (log_10_) scale.

The discrete Laplace method [2, 13] scrutinises the Y-STRs in the data set and identifies likely ancestral Y-STR profiles (haplotypes) and a likely distribution of the Y-STR profiles in the population from which the data set was sampled. This becomes more and more complicated and computer and time-consuming with increasing numbers of Y-STR loci. Thus, few Y-STR loci are preferred. More Y-STRs increase the polymorphism and the weight of evidence. However, due to the haplotypic nature of Y-STR profiles, the increase in polymorphism is dramatically decreasing when adding information from more and more Y-STR loci. The 17-Y-STR profile gave the highest weight of evidence with *LR*s above 10,000 in approximately 94% of the cases and above 100,000 in approximately 82% of the cases (Fig. 4 and Table 4). However, the 17-Y-STR profile had a slightly higher bias than the 12-Y-STR profile. However, the difference is so small that it has no practical consequence (Fig. 5). The median absolute deviation, *MAD*, defined in Eq. (4), was very similar among the three “kits”, and decreased rapidly similar to the bias (Fig. 5). For databases of sizes of at least 1,000, the *MAD* was smaller than 0.01.

The information of, e.g., a 12-Y-STR profile is more than 95% of that of a 21-Y-STR profile (Table 3) when estimated as suggested by [17]. The 12-Y-STR profile gave *LR*s above 10,000 in approximately 89% of the cases and above 100,000 in approximately 70% of the cases (Fig. 4 and Table 4), showing the value of the discrete Laplace method with also small numbers of Y-STRs. When the database includes approximately 1,000 Y-STR profiles, the difference between the *LR*s with and without adding the suspect’s Y-STR profile is neglectable for practical purposes (Fig. 5). The weight of evidence was over-estimated when the suspect’s Y-STR profile was not added to databases with less than 1,000 Y-STR profiles (Fig. 2 and Fig. 5).

No method takes into account that increased numbers of similar Y-STR profiles may be introduced in a population in case of, e.g., artificial insemination of a large number of women with semen from a single donor.

Naïve estimates of Y-STR profile probabilities like (*x* + 1)*/*(*n* + 1) or (*x* + 2)*/*(*n* + 2) (where *n* is the size of the database and *x* is the number of times the profile in question is present in the database) with a Y-STR database of size 1,000 will, in most cases, lead to *LR*s of 1,000 or 500. These figures are 100 to 1,000 times lower than those obtained with the discrete Laplace method. The discrete Laplace method is also superior to the naïve estimate for smaller Y-STR databases. A database with, e.g., 100 males will typically result in *LR*s of 100 or 50, while the discrete Laplace method results in *LR*s above 10,000 in approximately 94% of the cases with 17 Y-STR loci. However, if the database is small, the suspect’s matching Y-STR profile should be added to the Y-STR reference data to avoid over-estimating the weight of evidence, which results in time-consuming computations and the need for expert knowledge.

Most forensic genetic laboratories use commercial Y-STR kits with 17 or more loci. However, the loss of information is small when the information from, e.g., 12 Y-STR loci instead of 17 Y-STR loci is used for calculating the *LR*. For exclusion purposes, the information from all examined Y-STRs should, of course, be used to compare, e.g., the profiles from a scene of a crime and a suspect because of the increased power of discrimination of the additional loci that often include highly mutating loci.

### 4.3. Y-STR matches and close male relationship

Approximately 50% of the sharing of Yfiler Plus profiles with 25 Y-STR loci are within five meiotic steps from the proband (Fig. 7). This corresponds to, e.g., a second cousin relationship. For Y-STR kits with lower profile mutation rate, the fraction of sharing within five meiotic steps is decreased, and hence a larger fraction of the matches will be present in the general population. In many countries and societies, it is difficult to find other relevant information about male relatives beyond five meioses. Thus, the caseworkers handling criminal cases must be aware of the caution against close male relatives when the weight of evidence is presented as the *LR* including ‘a random man from the population’. Particularly, close male relatives to the suspect, like fathers, sons, uncles, nephews, cousins, second-degree cousins, etc., likely have Y-STR profiles similar to the suspect. Attempts to reduce the risk of similarity between close male relatives include using Y-STRs with high mutation rates. However, such Y-STRs are not optimal for use with the discrete Laplace method because the combined mutation rate of a Y-STR profile should be less than 5% to ensure that the matching individuals are found in the entire population and not only among the suspect’s close paternal relatives. However, this is a minor problem because most new kits with rapidly mutating Y-STRs will include the presently used Y-STRs with lower mutation rates. Thus, a higher discrimination power among close male relatives and a high *LR* among unrelated can be obtained.

### 4.4. Use of Y-STR typing in criminal cases

Y-STR typing is used in only a small proportion of criminal cases where it might be relevant. This is partly due to the limited weight of evidence reported when using the naïve method of *LR* calculation and the risk of Y-STR matches caused by close paternal relationships. Using the discrete Laplace method will increase the weight of evidence and most likely increase the use of Y-STR typing. A proportion of Y-STR profile matches between suspects and crime scene DNA are due to the fact that the crime scene DNA comes from a close relative to the suspect. In such cases, the information can be used for family investigations. Family searching in well-curated, up-to-date crime DNA databases with Y-STR and autosomal STR profiles is very useful and more cost-effective than forensic genealogy search based on Single Nucleotide Polymorphism (SNP) typing of, e.g., 600,000 SNPs and search in databases intended for other purposes [22, 23].

### 4.5. IT tool for the discrete Laplace method

We have developed a freely available IT tool, the R package `disclapmix` [13] available at https://cran.r-project.org/package=disclapmix, for creating the discrete Laplace model of a dataset and estimating the weight of evidence in criminal cases based on an already created discrete Laplace model. The R package contains vignettes with usage examples. The vignettes are available in the package or directly at https://mikldk.github.io/disclapmix/articles/. If a database includes approximately 1,000 Y-STR profiles or more, laboratories can establish the discrete Laplace model for the database and use the model for future work until the database is updated. When the discrete Laplace model is established, the *LR* calculation with the matching profile in a criminal case is performed in seconds, and the computations are easily documented.

### 4.6. Conclusion

We have shown that the discrete Laplace method offers a reliable way of estimating the weight of evidence of matching Y-STR profiles in criminal cases. The *LR* can be calculated in seconds with databases with 17 STRs from at least 1,000 males once the discrete Laplace model of the population, from which the database was sampled, has been established. Approximately 50% of the Y-STR matches with the Yfiler Plus kit (25 Y-STR loci) are between male relatives within a genetic distance of five meioses. We offer freely available IT tools and computer code for (1) discrete Laplace modelling of individual Y-STR databases and (2) fast calculation of *LR*s of matching Y-STR profiles.

